# Experimental variables that impact outcomes in *Caenorhabditis elegans* aging stress response

**DOI:** 10.1101/2024.01.09.574889

**Authors:** Bradford Hull, Isabella M. Irby, Kayla M. Miller, Ally Anderson, Emily A. Gardea, George L. Sutphin

**Affiliations:** University of Arizona

**Author notes:** Corresponding author: George L. Sutphin.

## Abstract

Cellular stress is a fundamental component of age-associated disease. Cells encounter various forms of stress – oxidative stress, protein misfolding, DNA damage, etc. – and respond by activating specific, well-defined stress response pathways. As we age, the burden of stress and resulting damage increases while our cells’ ability to deal with the consequences becomes diminished due to dysregulation of cellular stress response pathways. Many interventions that extend lifespan activate one or more stress response pathways or allow cells to maintain normal stress response later in life. The nematode *Caenorhabditis elegans* is a commonly used model for both aging and stress response research. As such, stress response experiments are regularly conducted as part of studies focused on mechanisms of aging in *C. elegans*. However, experimental design across experiments in the field are highly variable, including stressor dose, age at exposure, culture type (liquid vs. solid), bacterial strain used as a food source, and environmental temperature. These differences can result in different experimental outcomes, making comparison of results between studies challenging. Here we evaluate several experimental variables that are variable in the published literature and find that each can meaningfully alter experimental outcomes for multiple stressors. Our goal is to raise awareness of the issue of experimental variability within the field and suggest a standardized experimental design to serve as a set of guidelines for future experiments. By adopting these guidelines as a starting point, and explicitly noting differences in specific experiments, we aim to promote rigor and reproducibility, ultimately fostering more interpretable and translatable outcomes in geroscience research.

## Introduction

Stress is intricately linked to aging. Cells and organisms are continuously challenged with various forms of stress—reactive oxygen species, changes in temperature, misfolded proteins—and in response activate a diverse set of molecular pathways to help mitigate or repair the resulting damage. As we age, our cells experience an increasing burden of stress, while the molecular response pathways become dysregulated. Understanding the mechanisms linking stress and stress responses to aging and targeting these mechanisms to extend healthy lifespan and combat age-associated disease is an ongoing focus of aging research.

The roundworm *Caenorhabditis elegans* is a long-standing and widely used model system for both aging [1, 2] and stress response [3, 4]. *C. elegans* are used extensively in both aging and stress response due to desirable characteristics such as their short lifespan, inexpensive culture, easy delivery of stressors, and a plethora of well-developed tools, such as RNAi and fluorescent stress-response reporters. Many *C. elegans* stress response pathways are highly conserved when compared to their human counterparts. Induction and activation of stress response pathways in response to physiological stress has been well documented in nematodes across many types of stress including UV exposure, oxidative stress, hypoxia, hyperosmolarity, heavy metal exposure, and heat shock.

Stress response experiments are typically performed by establishing a synchronized population (via bleaching or other methods), exposing worms to a chemical or environmental stress, and monitoring survival, health, or some other molecular output at one or more subsequent timepoints. However, these experiments in the *C. elegans* stress response field lack standardization, creating results that are often difficult to interpret in the context of similar research. Experimental variables like stressor dose (e.g., acute vs. chronic), culture conditions, age at stress exposure, strain of bacterial food source, and method of stressor delivery can meaningfully impact the stressor dose needed to achieve a specific health outcome, activation of stress response pathways, and longevity of the organism [5–7], which can in turn significantly affect both experimental outcomes and biological interpretation of a study. This is particularly important for building an accurate understanding of biology of stress response by examining the body of evidence from experiments across different projects and laboratories. For experiments to be directly comparable, they need to be completed under similar conditions. Drawing conclusions about the impact of a stress and the underlying molecular response of the organism should take into account experimental differences between experiments and studies, and the level of confidence placed in these conclusions moderated according to our understanding of the influence of relevant experimental variables on outcomes.

Some experimental variables have specific considerations that are relevant to understanding the impact of stress and stress response in the context of aging. For example, many studies evaluate the impact of a stressor by measuring survival or health of *C. elegans* challenged at the L4 larval stage, while animals are at a stage of rapid cellular growth. Many stressors cause *C. elegans* to slow or arrest growth, particularly when applied at acutely toxic doses. L4 animals are still undergoing a state of transition, with some tissues (e.g., the vulva [8]) undergoing substantial morphological and transcriptional changes. Molecular responses to external stimuli, including stressors like heat [9], are distinct from those of mature adult animals, and interpretation of outcomes assessed at the L4 stage should take this into consideration and not assume that the molecular responses during development will be identical to those during adulthood. Similar considerations should be made for young vs. aging animals [6].

Here we discuss experimental variables that frequently vary between labs and publications in the context of aging stress response research and explore how altering these variables can affect organismal survival, health, and activation stress response pathways. We present data using copper, dithiothreitol (DTT), and, in select instances, sodium as examples of exogenous stressors with different mechanisms of action that elicit distinct responses to changes in different experimental variables. Finally, we propose a set of standard experimental conditions as a reasonable starting point for experiments designed to probe mechanisms of stress response in aging *C. elegans*. These guidelines will allow nematode stress response research to be standardized, which will aid researchers in interpreting findings from different labs and make future experiments more translatable, from both lab to lab and bench to bedside.

## Methods

### Strains

All experiments used wild type N2 worms, originally obtained from Dr. Mat Kaeberlein (University of Washington, Seatle, WA, USA).

### Solid media and culture

*C. elegans* were cultured on solid nematode growth media seeded with *E. coli* bacteria as previously described [10]. Briefly, all solid media experiments were conducted on NGM containing agar and 25 μg/mL carbenicillin. Experiments using OP50 excluded carbenicillin. All plates were seeded with live *E. coli* (HT115), except where stated that OP50 was used. Worms were age-synchronized via bleach preparation and transferred at L4 larval stage to plates containing 50 μM 5-fluorodeoxyuridine (FUdR) to prevent reproduction [10]. Stressors were introduced by adding either copper sulfate (CuSO_4_, Fisher Chemical, catalog number C493-500) or dithiothreitol (DTT, FisherBiotech, catalog number BP182-25) to NGM media during preparation.

### Lifespan analysis

Lifespan experiments were completed as previously described [10]. Worms were placed on NGM plates containing FUdR and the desired stressor on day 2 of adulthood except where otherwise noted. Each animal was examined every 2-3 days by nose- and tail-prodding with a platinum wire pick. A worm was considered dead if they failed to react to prodding and then removed from the plate. Worms showing vulva rupture were included in analyses, while those that left the surface of the plate were excluded. Rims of plates were coated with 10 mg/mL palmitic acid in 100% EtOH to discourage fleeing. Each experiment was measured in three independent repeats. P-values for statistical comparison of lifespan between conditions were calculated using the log-rank test (survdiff function in the R “survival” package) with Holm multiple test correction.

### Body size measurements

Worms were synchronized and raised as described above. All worms were transferred to NGM plates containing FUdR at the L4 larval stage. When worms reached the appropriate age (L4, day 2, or day 4 of adulthood), they were transferred to control plates or plates containing the indicated stressor. Pictures were taken at day 8 of adulthood and body size measured using the measurement tool in arivis Vision4D software (version 4.1.1, https://www.arivis.com/products/pro). Settings were consistent across all images.

### Brood size

To measure brood size, 10 worms at the L4 larval stage were placed on individual NGM plates without FUdR and allowed to lay eggs. Worms were transferred to new plates every 24 hours until they stopped laying eggs. Plates were incubated for two days to allow eggs to hatch and the number of progeny was then counted. Two-tailed Welch’s t-tests were performed to determine significance between treatment and control groups.

## Results

For experiments discussed in this paper, the baseline conditions are generally as follows: day 2 adults on NGM agar plates spoted with *E. coli* HT115 and with the stressor dissolved in the media. Each experiment discussed below tests each aspect of this general experimental setup and argues a case for why this choice was proposed as the standard.

### Age of Exposure

The age at which worms are exposed to a stressor can often have a significant impact in their response to that stressor [6]. While there are more or less common ages that are used (L4, early adult), these ages are highly variable between studies and often even within the same paper [11–13]. In some cases, the age at which worms are challenged may be problematic for the question being addressed (e.g., assessing stress response of larval animals in the context of an aging intervention) or stated imprecisely (e.g., “early adult” instead of “day 1 of adulthood”). Given the brief life cycle of *C. elegans*, a difference of even a day or two in age of exposure can yield worms with distinct morphology [8] and stress resistance [6]. Interpreting data from animals exposed to stress during larval stages should consider possible impacts on animal development that may affect phenotypes much later in life. Since somatic cells in the worms are still mitotic until the L4 larval stage, earlier exposure to an aversive intervention could potentially disrupt cell division or any of the other growth processes that are still occurring prior to adulthood. During the larval stages, worms are also still undergoing cuticle development and gonadogenesis, both processes which may affect a worm’s ability to respond to stress. L4 worms that have acquired all of their somatic cells are still in a phase of rapid growth even if the somatic cells are not dividing, and exposure to stress can potentially affect growth during this period. In contrast, a worm that is allowed to reach old age before exposure to stress will tend to have lower stress tolerance. Worms that have been given time to develop past larval stages often have higher tolerances for stressors than larva, with stress resistance peaking in early adulthood [6]. Examining worms at the peak of their stress resistance has advantages in terms of reducing variability and simplifying interpretation, but also disadvantages if the toxic dose of a stressor at this stage exceeds the solubility the media, or if a chemical stressor is particularly expensive. In agreement with Dues et al. [6], we find that worms exhibit greater stress tolerance to multiple forms of stress in early adulthood when compared to the commonly used exposure at L4 under copper sulfate and sodium stress (**Figure 1A-D)**, but not DTT stress **(Figure 1E,F**). We also see that exposure to stress at L4 results in worms that are smaller in size compared to those exposed at day 2 of adulthood, but exposure to stress at L4 does not affect brood size, suggesting that these select stressors affect growth but not reproduction (**Figure 2**).

**Figure 1.**
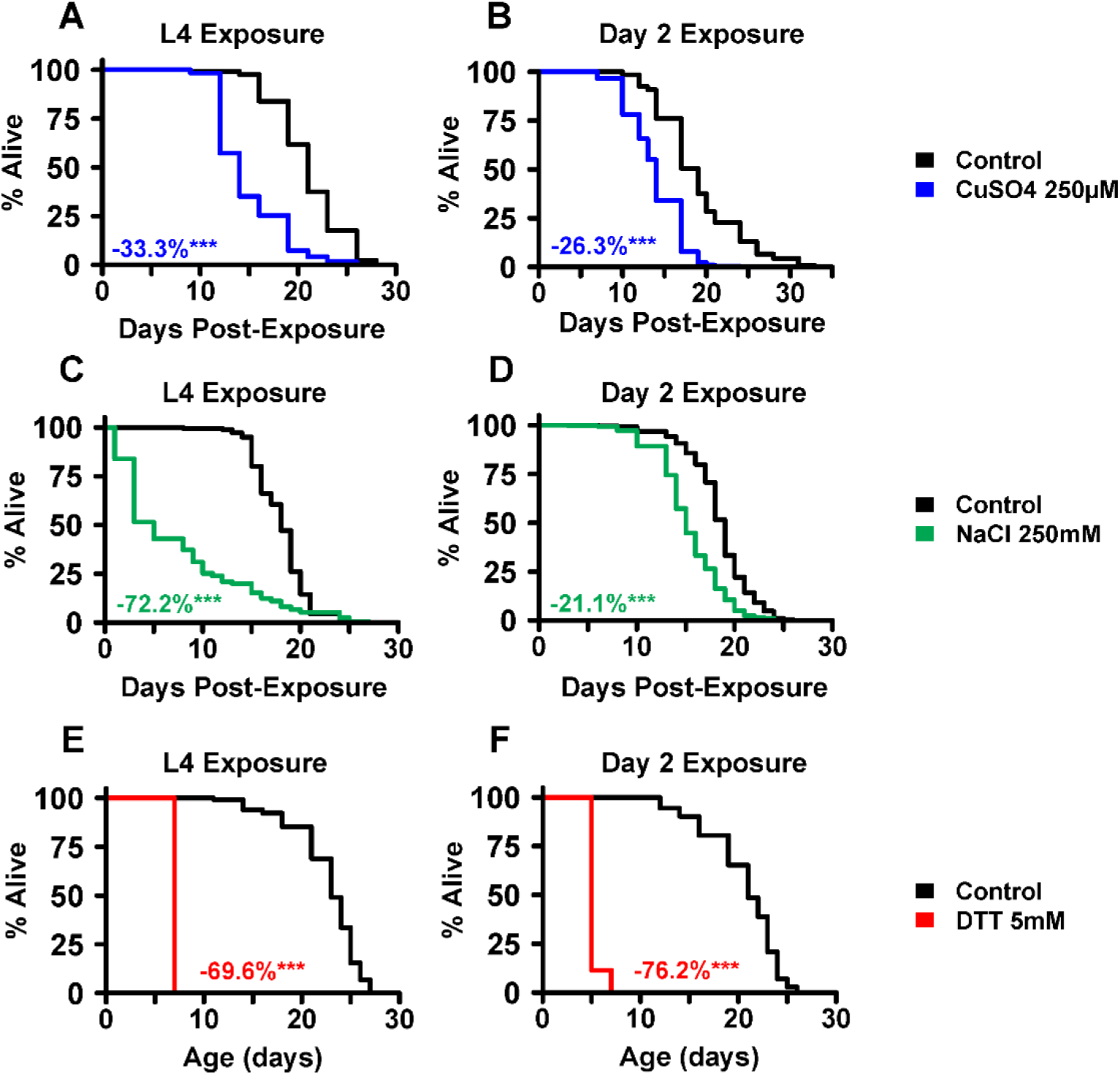
Age at time of stress exposure affects toxicity and development. Exposure to stress is significantly more toxic when delivered at the L4 larval stage compared to day 2 of adulthood for both CuSO_4_ and sodium (NaCl) **(A-D)**. However, DTT exposure did not show the same marked difference **(E**,**F)**. Significance compared to the controls is noted with * p < 0.05, ** p < 0.01, *** p < 0.001.

**Figure 2.**
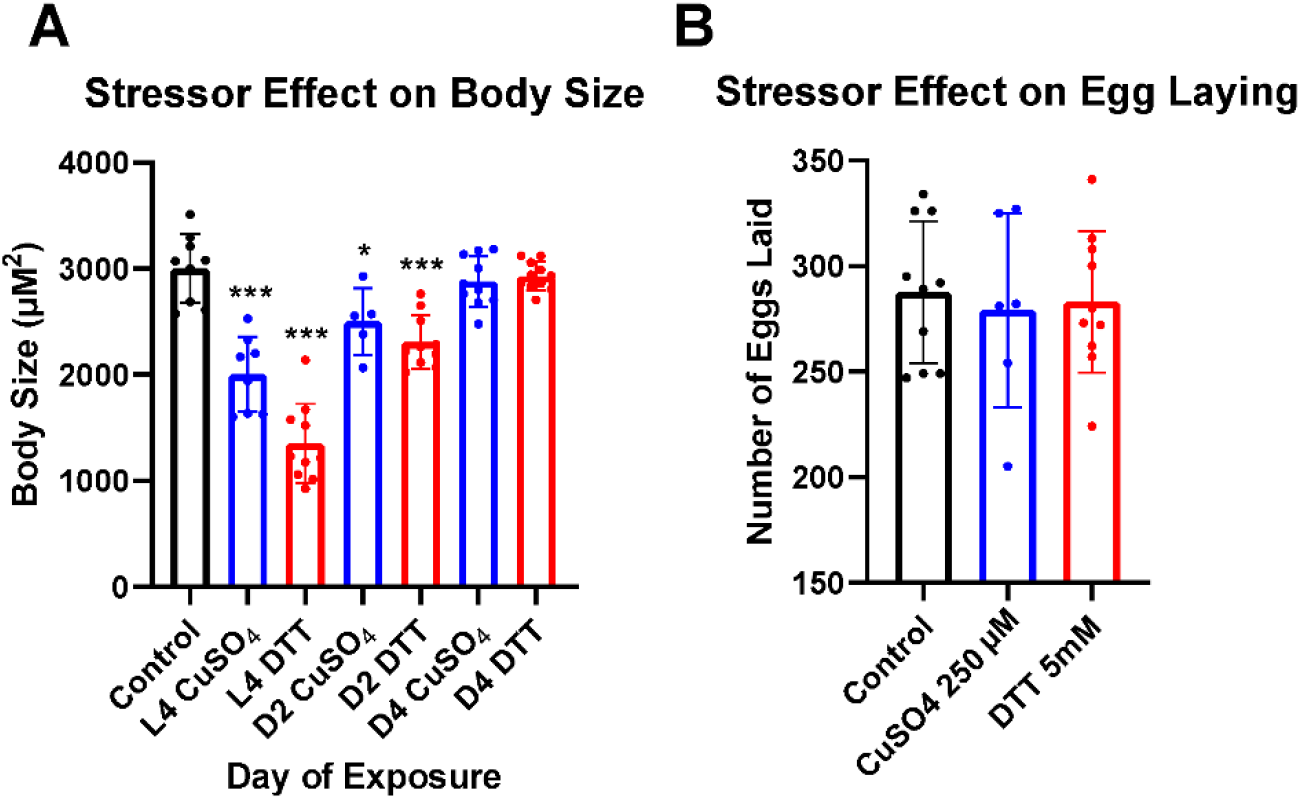
Stress affects body size but not brood size. **A)** Exposure to CuSO_4_ or DTT starting at the L4 larval stage results in a smaller final body size. Exposure at day 2 of adulthood also results in a smaller body size, but to a lesser extent. Body size was measured for all worms at day 8 of adulthood. **B)** Exposure of L4 larval worms to CuSO_4_ or DTT stress does not significantly affect brood size. Significance compared to the controls is noted with * p < 0.05, ** p < 0.01, *** p < 0.001.

### Method of Exposure

The method of exposure by which a drug is introduced to nematodes is more consistent across the literature, with most publications adding the drug into the media before plates are poured [14–16]. However, there are some situations in which it may be advisable to introduce the drug via a different method, such as when the chemical is sensitive to heat, exhibits poor solubility in water-based media, or is expensive. In such cases, drugs can be delivered topically either prior to adding food or mixed with *E. coli* food [17–19]. Stressors added topically are generally considered to diffuse evenly across the media, though this will depend on several factors including the solubility of the drug and potential uptake of the drug by the bacteria. Here we discuss and compare the effects of stressors when introduced via media or topical addition, using copper sulfate and DTT stress as examples.

We find that lifespan of worms exposed to 250 μM copper sulfate experience a 22% lifespan reduction whether the copper is added to the media or to the bacteria (**Figure 3A**). This suggests that copper is able to diffuse evenly throughout the plate when initially mixed with the bacteria so that the final effective dose experienced by the worms is the same. In contrast, 5mM DTT reduces lifespan by 52% when added to the media, but by 69% when added to the bacteria (**Figure 3B**). Because DTT is more toxic for worms when added by mixing with the bacterial food, it is likely that DTT either does not diffuse well throughout the media, is taken up and concentrated in the bacteria, or has an indirect impact on survival by altering the behavior of the bacteria.

**Figure 3.**
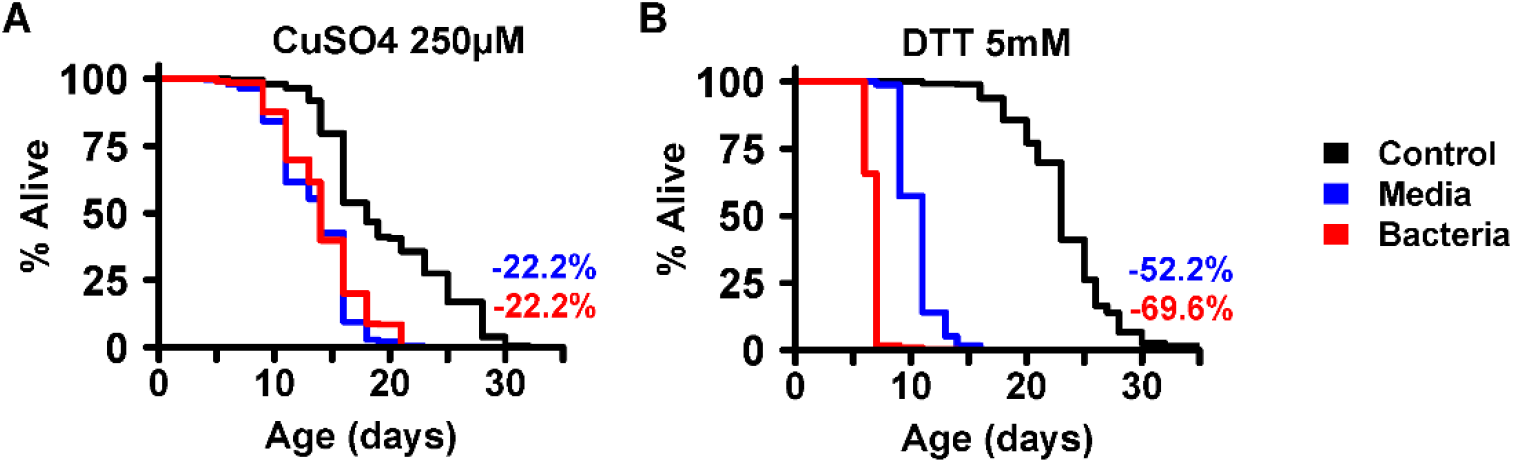
Stressor delivery can significantly affect survival. **A**) The toxicity of CuSO_4_ stress was similar when delivered via addition to the growth media or the bacterial food. **B**) DTT stress was significantly more toxic to worms when delivered via bacterial food rather than through the growth media. Significance compared to the controls is noted with * p < 0.05, ** p < 0.01, *** p < 0.001.

### Bacteria

The species and strain of bacteria used to feed nematodes varies substantially across the *C. elegans* stress response literature. While many experiments use either the *E. coli* strains OP50 (often considered the default strain in *C. elegans* research) or HT115 (the parental strain for both RNAi feeding libraries [20, 21]), there are a plethora of other strains used including *E. coli* P90C and HB101 [22, 23]. Though type of food is often considered less important than other experimental variables, the species and even the strain of bacterial food can significantly affect how worms respond to aversive interventions [15, 24, 25]. Many other species of bacteria have also been found to influence stress tolerance, usually when compared to *E. coli* OP50 [24].

We compared the effects of copper sulfate and DTT on worm lifespan when fed either *E. coli* OP50 or HT115 and found significant differences in *C. elegans* stress tolerance. When fed a diet of *E. coli* OP50, worms exposed to copper sulfate lived 9% shorter than those fed a diet of HT115 (**Figure 4A,B**). When exposed to DTT, OP50-fed worms lived 32% longer than HT115-fed worms (**Figure 4C,D**). These results corroborate the work discussed above and suggest a significant role of bacterial food in *C. elegans* stress response research.

**Figure 4.**
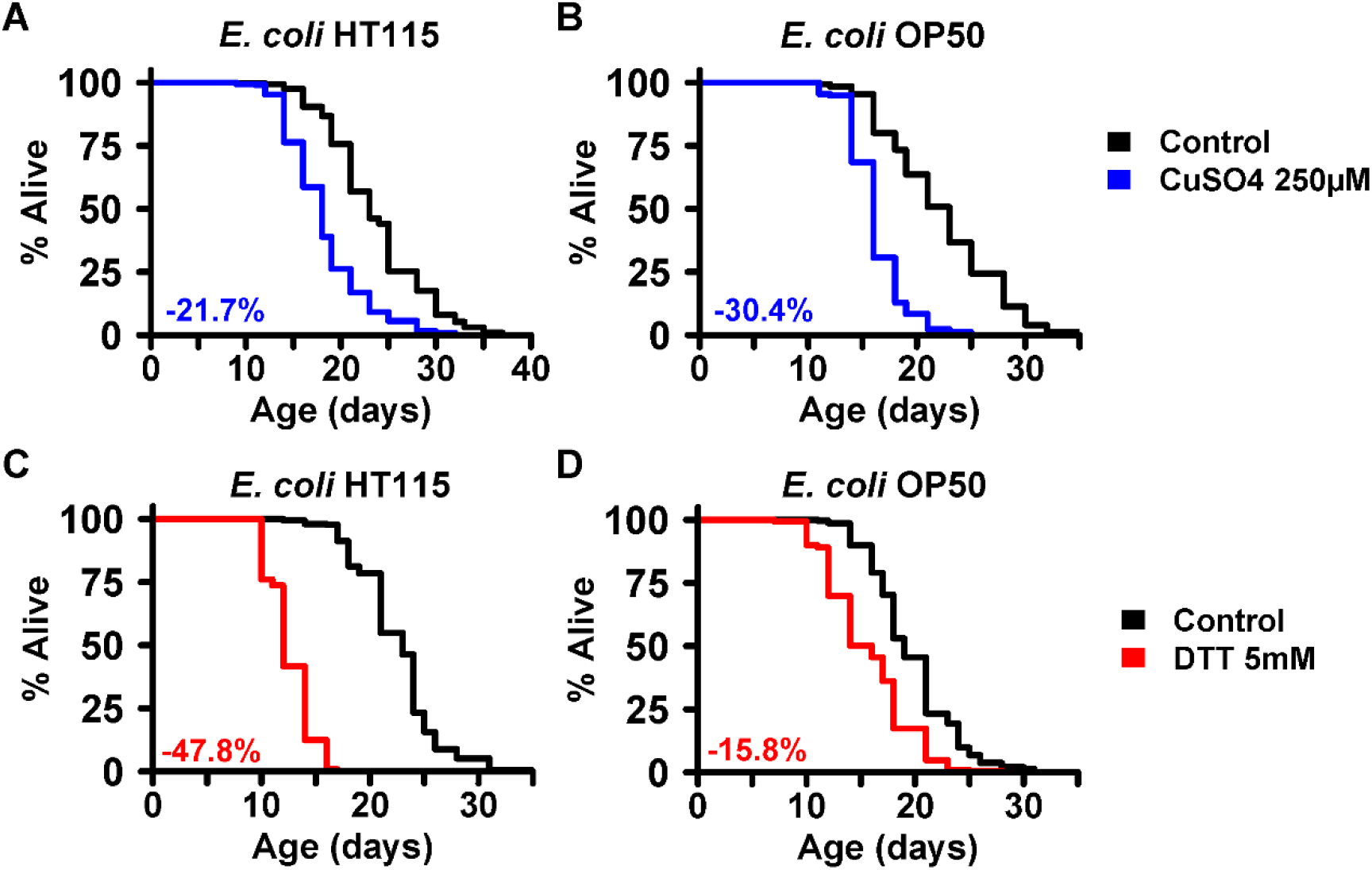
Bacterial food type significantly affects stressor toxicity to *C. elegans*. *C. elegans* experienced a modestly reduced toxicity from CuSO_4_ stress when fed *E. coli* HT115 bacteria (**A**) as compared to *E. coli* OP50 (**B**). *C. elegans* experienced a greatly enhanced toxicity from DTT stress when fed *E. coli* HT115 bacteria (**C**) as compared to *E. coli* OP50 (**D**). Significance compared to the controls is noted with * p < 0.05, ** p < 0.01, *** p < 0.001.

## Discussion

Here we evaluate and discuss multiple variables relevant to *C. elegans* stress response experiments and show how altering these variables can affect experimental outcomes. We first discussed the age of stressor exposure and showed that early adulthood is the optimal age over the L4 larval stage for exposure in a general stress response experiment. At this exposure age, most of the concerns about interpreting experiments where larval worms are challenged discussed above are avoided. Early adulthood is also thought to be the age of maximum stress resistance [6], which makes interpretation simpler and allows for easier comparison between studies. Additionally, we suggest the second day of adulthood because it allows for pre-exposure treatments with stressors (i.e. hormesis experiments) during the first day of adulthood while still avoiding larval exposure and keeping the time of exposure to the main stressor consistent.

We next considered the method by which stressors are delivered to worms and propose that the standard of stressor delivery be via agar media. The results of **Figure 3** show that the method of delivery in a stress response experiment can be important when comparing results between experiments, and that certain stressors may behave differently from others regarding diffusion, bacterial uptake, modification of the stressor by the bacteria, or changes to bacterial behavior. Most experiments in the literature deliver stressors via media and we propose to keep this as the standard. Although certain stressors may require topical delivery or another method due to sensitivity to heat, poor solubility in water-based media, or high cost, but outside of those fringe cases delivery via media allows for a direct comparison to most published studies to date and ensures stressor homogeneity across the plate.

We next examined the type of bacteria used as a food source during stress response experiments. We showed that different strains of bacteria can have significantly different impacts on survival of worms exposed to aversive interventions. These differences could be due to factors like uptake efficiency of the stressor by the bacteria, direct health effects of the bacteria on the worm (e.g., B12 deficiency, different levels of pathogenicity [26, 27]), or other possibilities. We propose a standard of using *E. coli* HT115 for stress response experiments in *C. elegans*. Beyond the reasons listed above, many stress response experiments are also done in conjunction with RNAi, which usually uses HT115 with a plasmid containing the RNAi gene of interest. Therefore, conducting non-RNAi stress response experiments with HT115 will allow them to be more directly comparable to RNAi experiments in the field. To note, an RNAi-compatible OP50 strain is now available, but that genome-wide feeding libraries are only available for HT115 [28]. It is worth noting that we have not discussed bacterial food pathogenesis in detail here. It is known that bacterial food can infect worms and impact survival [29, 30]. There are methods for working around this pathogenesis (UV or heat-killed bacteria, for example), but they come with their own set of limitations [10]. UV killing of bacteria is low throughput and inconsistent [31], and heat killing destroys essential nutrients for worms in the plate and can cause early developmental arrest [32]. Another potential solution is with paraformaldehyde killing, which the Leiser lab has shown to be effective and consistent [33].

Stress response experiments are a core component of the aging field, as physiological stress is a frequently experienced condition that contributes to aging and age-associated disease. A current major issue in the field is that it is often difficult to compare experiments between studies due to lack of standardization in methodology. We propose the following set of experimental conditions as a standard starting point when studying stress response in the context of aging:

- Exposure to stress at day two of adulthood
- Dissolve stressor in media for delivery
- *E. coli* HT115 for bacterial food

We intend these standards to be a framework for future experiments that can be used as a starting point and recognize that the goals of specific experiments will require deviation. By adopting the standard as a starting point in these cases, the deviation from the standard can be justified in the context of the experiment and explicitly noted in the experimental methods. In summary, adoption of a set of standards will not only improve the rigor and reproducibility of experiments in the stress response in aging field, but will also lead to faster and more robust translational interventions for geroscience overall.

## Acknowledgements

This work was supported by NIH/NIGMS award R35GM133588 to G.L.S and NIH/NIGMS award T32GM136536.

